# The effects of awareness of the perturbation during motor adaptation on hand localization

**DOI:** 10.1101/410753

**Authors:** Shanaathanan Modchalingam, Chad Michael Vachon, Bernard Marius ’t Hart, Denise Y. P. Henriques

**Affiliations:** Kinesiology and Health Science / Center for Vision Research, York University, Toronto, Ontario, Canada

## Abstract

Explicit awareness of a task is often evoked during rehabilitation and sports training with the intention of accelerating learning and improving performance. However, the effects of awareness of perturbations on the resulting sensory and motor changes produced during motor learning are not well understood. Here, we use explicit instructions as well as large rotation sizes to generate awareness of the perturbation during a visuomotor rotation task and test the resulting changes in both perceived and predicted sensory consequences as well as implicit motor changes.

We split participants into 4 groups which differ in both magnitude of the rotation (either 30° or 60°) during adaptation, and whether they receive a strategy to counter the rotation or not prior to adaptation. Performance benefits of explicit instruction are largest during early adaptation but continued to lead to improved performance through 90 trials of training. We show that with either instruction, or with large perturbations, participants become aware of countering the rotation. However, we find a base amount of implicit learning, with equal magnitudes, across all groups, even when asked to exclude any strategies while reaching with no visual feedback of the hand.

Participants also estimate the location of the unseen hand when it is moved by the robot (passive localization) and when they generate their own movement (active localization) following adaptation. These learning-induced shifts in estimates of hand position reflect both proprioceptive recalibration and updates in the predicted consequences of movements. We find that these estimates of felt hand position, which reflect updates in both proprioception and efference based estimates of hand position, shift significantly for all groups and were not modulated by either instruction or perturbation size.

Our results indicate that not all processes of motor learning benefit from an explicit awareness of the task. Particularly, proprioceptive recalibration and the updating of predicted sensory consequences are largely implicit processes.

## INTRODUCTION

In many motor learning tasks, participants learn to compensate for a systematic perturbation applied to an effector such as their reaching hand. The perturbation may include changes in the dynamics involved in producing movements (Shadmehr & Mussa-Ivaldi, 1994; Smith, Ghazizadeh, & Shadmehr, 2006), inertial loads attached to moving body parts (Krakauer, Ghilardi, & Ghez, 1999) or changes in the visual feedback associated with the movements (Krakauer, Pine, Ghilardi, & Ghez, 2000). Learning during such paradigms is initially rapid, with eventual decrease in the rate of improvements of compensations until performance asymptotes near pre-perturbation baseline levels; this pattern of improvement constitutes a typical learning curve. Learning to return to baseline levels of performance in presence of a perturbation is known as adaptation.

Adaptation involves both implicit, and explicit, conscious processes. Typically, adaptation to small perturbations is thought to be implicit, where there is a change in an internal forward model due to sensory prediction errors brought about by the perturbation (Mazzoni & Krakauer, 2006; Tseng, Diedrichsen, Krakauer, Shadmehr, & Bastian, 2007). The implicit processes associated with motor adaptation may be gleaned by measuring the motor aftereffects; the continued change in performance even in the absence of the perturbation (Krakauer et al., 1999, 2000; Redding & Wallace, 2000). However, it has been suggested that people also use cognitive strategies to aid in the adaptation process, especially during early adaptation (McDougle, Bond, & Taylor, 2015; McDougle, Ivry, & Taylor, 2016). Furthermore, there are cases such as with large rotations (Bond & Taylor, 2015; Hegele & Heuer, 2013; Neville & Cressman, 2018; Werner et al., 2015) or with prior instructions (Benson, Anguera, & Seidler, 2011; Heuer & Hegele, 2008; Neville & Cressman, 2018; Werner et al., 2015) where participants continue to employ explicit strategies well after adapting to the rotation. Here we attempt to evoke awareness during the learning process.

When we exert movements, such as reaches, we generate predictions of the consequences of the initiated motor command based on a copy of the motor command. Introducing perturbations causes a consistent discrepancy between the predicted consequences of a movement, and the actual experienced consequences; i.e. the sensory feedback received during the movement. This discrepancy may then be used to improve future movements by changing the mapping between motor commands and their predicted outcomes (Bastian, 2008; Berniker & Kording, 2008). When the perturbation is a change in visual feedback, where visual consequences of reaches are manipulated, there is additional discrepancy between different modalities of sensory feedback. Specifically, our estimates of hand position based on proprioceptive inputs are decoupled from those based on visual input. This discrepancy results in a remapping of the relationship between proprioceptive feedback and the perceived location of the hand in space; a process known as proprioceptive recalibration (Cressman & Henriques, 2009; Cressman, Salomonczyk, & Henriques, 2010; Ruttle, Cressman, ’t Hart, & Henriques, 2016; Salomonczyk, Cressman, & Henriques, 2011). Our ultimate perceived hand position is informed by both the predicted consequences of an initiated movement, and the received sensory feedback (’t Hart & Henriques, 2016).

The effects of awareness when adapting to visuomotor rotation on corollaries of motor learning, such as the changes in predicted consequences of movements and proprioceptive recalibration are poorly understood. Here, we evoke explicit strategy use with instructions and large rotation sizes and examine the effects on changes in prediction and proprioception. We hypothesize that when people are aware of the nature of the perturbation, they will exhibit lower amounts of recalibration of felt hand position, as the systematic errors experienced during training will be attributed to external sources rather than the body. Thus, we predict that changes related to the perception of where the body is located will be reduced. We also predict that a 60° visuomotor perturbation is sufficiently large to lead to explicit awareness of the perturbation without the requirement of explicit instructions whereas a 30° perturbation is not. Thus, when participants experience a 60° rotation, we expect them to show learning resembling instructed groups and an ability to modulate their strategy use when cued, even when they are not provided with instructions on the nature of the perturbation. By using this method of evoking “awareness”, we will test the implication that explicitly-driving learning can lead to changes in afferent and efferent-based hand localization.

## METHODS

### Participants

Eighty right-handed participants from York University took part in the experiment. All participants gave prior, written, informed consent and participation was voluntary. The procedures used in this study were approved by the York Human Participants Review Sub-committee. All participants reported having normal or corrected to normal vision.

The participants were separated into 4 groups. About half the participants adapted to a 30° visuomotor rotation and the other half to a 60° rotation. To ensure awareness during training, these two groups were further split by giving explicit instructions on the nature of the perturbation, and strategies to counter the rotation, to half of the participants who experienced each rotation size. Thus, there were four groups: non-instructed 30° -- NI30 (n = 20, 14 female), instructed 30° -- I30 (n = 21, 13 female), non-instructed 60° -- NI60 (n = 19, 14 female) and instructed 60° -- I60 (n = 24, 18 female). We visually inspected all ‘Reach to Target’, ‘Reach with No Cursor’, and ‘Localization’ tasks to ensure participants followed their given instructions. We excluded 8 participants who failed to complete the tasks as instructed.

### General set-up

Participants were seated on a height-adjustable chair in front of an apparatus (Fig 1C) which included a downward facing computer screen (Samsung 510 N, 60 Hz) located 28cm above a 2-joint robot manipulandum (Interactive Motion Technologies Inc., Cambridge, MA, USA). The chair’s height was adjusted until the participant could manipulate and reach with the robot handle comfortably while viewing the entire display on a reflective surface located 14cm above the robot manipulandum. Participants were then asked to grip a vertical handle on the manipulandum with their right hand. A thick black cloth was draped over participants’ right shoulder and arm which occluded the view of their right arm. The experiment was performed in the dark to limit peripheral vision of their right arm. The participants’ left hands were illuminated by small lamps. Visual stimuli were presented on the reflective surface using the downward facing screen. The reflected image appeared on the same horizontal plane as their right thumb.

**Fig 1:**
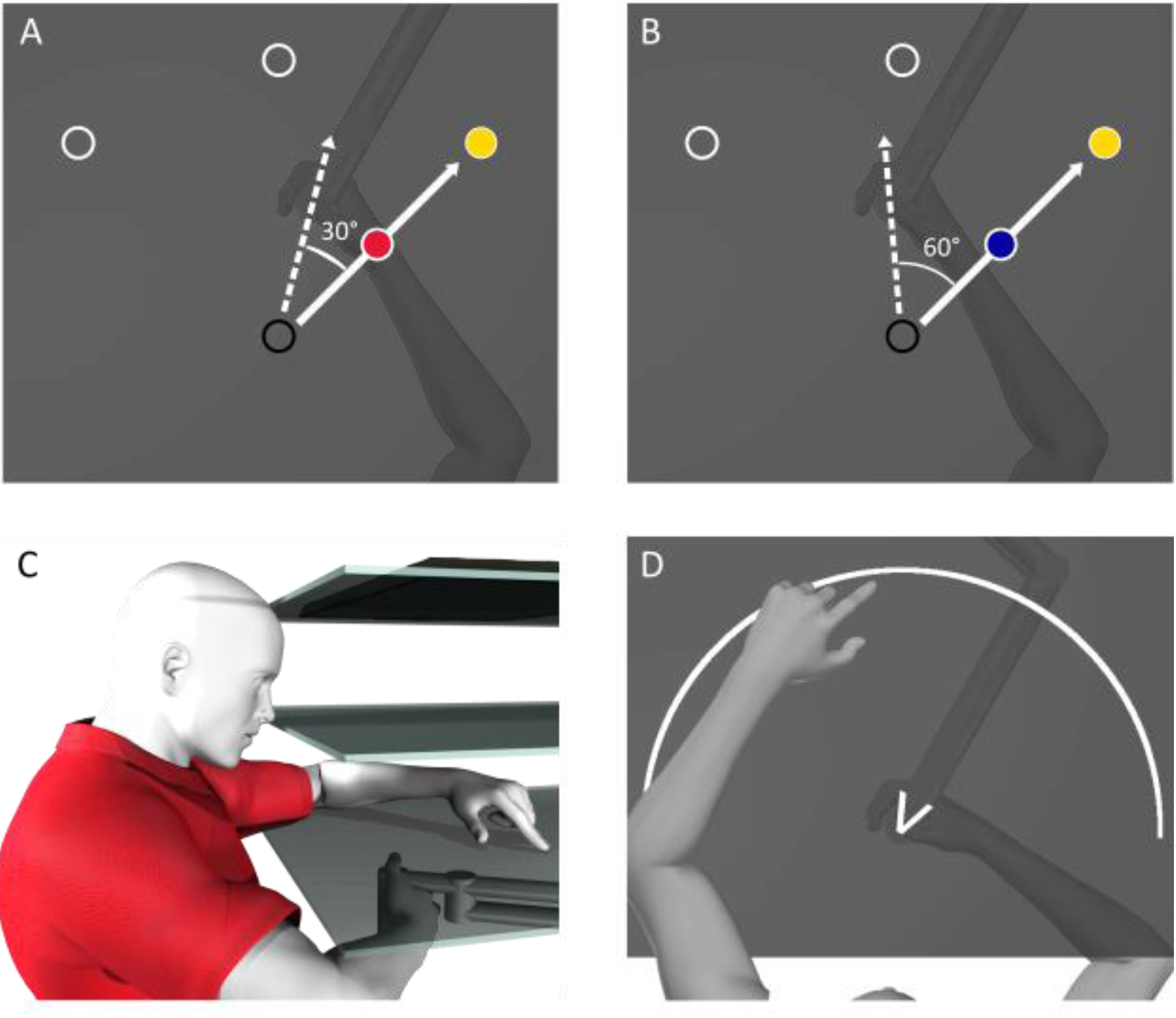
Experimental Setup and Procedure. A+B) During ‘Reach to Target’ tasks in the ‘Rotated’ session, the position of a cursor representing the hand (red and blue circle) was rotated 30° (A) or 60° (B) CW during rotated training tasks. Participants attempted to one of the yellow target as quickly and as straight as possible. C) Participants gripped a robot manipulandum located below a touchscreen (bottom surface) while looking at a reflective screen (middle surface). The reflected visual stimuli were projected from a monitor (top surface) located above the reflective screen. D) Participants used their visible left hand to indicate where they crossed an arc with their unseen right hand during localization tasks.

During ‘Reach to Target’ and ‘Reach with No Cursor’ tasks (Fig 2), participants made reaching movements to one of three possible visual targets (a circular disk 1 cm in diameter) that were situated 10 cm away from the starting position. For each trial, this target was located either straight in front of or at a 45° angle CW and CCW to the starting position (Fig 1A+B). During some tasks the participants used their visible left hand to indicate, on a touchscreen located 2cm above the manipulandum, the perceived position of their unseen right hand. Between tasks all visual stimuli were removed, and participants were instructed to return the robot handle to a starting position located 20cm in front of the body midline.

**Fig 2:**
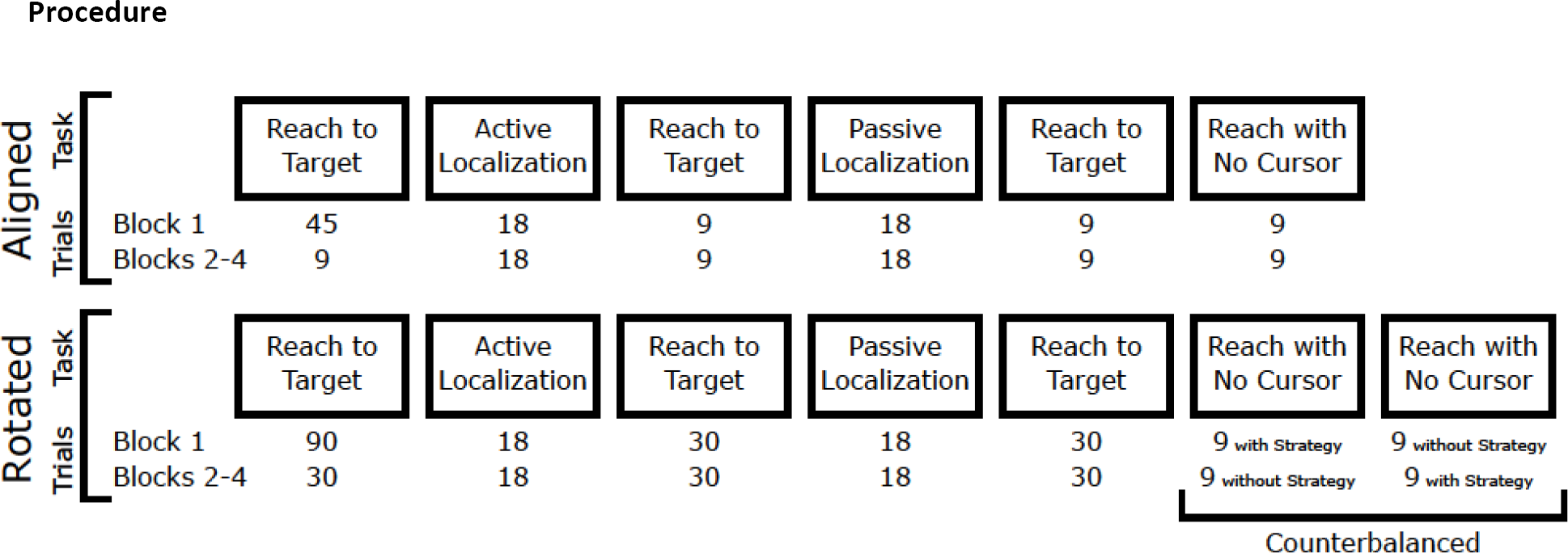
Sequence of Tasks. All four groups followed the same sequence of tasks. Participants completed four blocks of the aligned session of the experiment and took a mandatory 10-minute break before moving onto the rotated session. The “Reach to Target” tasks in the Rotated session either included a 30° or a 60° CW visuomotor rotation, depending which group the participant belonged to.

### Procedure

All participants completed ‘Reach to Target’ tasks followed by ‘Localization’ and ‘Reach with No Cursor’ tasks (Fig 2 and described in detail below). During ‘Reach to Target’ tasks in the ‘Aligned’ session of the experiment (top row of Fig 2), the location of a cursor representing the participant’s unseen hand was aligned with the real position of the participants’ unseen right hand. These tasks in the ‘Aligned’ session both familiarized the subject to the experiment and served as baseline data to which other data could be compared. After ‘Localization’ and ‘Reach with No Cursor’ tasks, participants completed top-up ‘Reach to Target’ trials. After completing blocks 1-4 in the ‘Aligned’ session, participants took a mandatory 10-minute break. In ‘Reach to Target’ tasks after the break, the motion of the cursor representing the unseen right hand was rotated about the starting position. The magnitude of the rotation was 30° CW (Fig 1A) for the NI30 and I30 groups, and 60° CW (Fig 1B), for the NI60 and I60 groups. During the mandatory break, the ‘Instructed’ groups were informed of the rotation and were provided with a strategy to counteract the rotation so that they could still move the cursor in a straight line to their targets. Specifically, they were instructed to visualize the starting position as being at the center of the clock face. They were told reaching towards a specific number on the clock would result in the cursor heading toward either the next number over (for the 30° rotation), or the second number over (for the 60° rotation) in the clockwise direction. The implicit groups were informed that the reach training tasks would feel different after the break, but no strategy or details were provided. Both groups were told to keep the strategy they were using to achieve straight reaches in mind, as they would be asked to recall and either use or not use the strategy during some ‘Reach with No Cursor’ trials.

#### Reach to Target tasks

Participants received visual feedback of their hand position via a continuously displayed green cursor. This circular green cursor, 1 cm in diameter, represented the location of their unseen right thumb. After participants placed their hand at the starting position for 300 ms, the target appeared. Participants were told to reach to the target as quickly and accurately as possible. A reach trial would end when the center of the hand cursor overlapped the target within 0.5 cm of the target’s center. When the reach was completed, both the target and the cursor disappeared. Participants then moved their hand back toward the starting position along a robot-constrained straight path, which was generated by a perpendicular resistance force of 2 N/(mm/s) and a viscous damping of 5 N/(mm/s). 45 trials of the ‘Reach to Target’ task were done at the start of the experiment, during block 1, and 9 trials were done during all other ‘Reach to Target’ tasks during the ‘Aligned’ session of the experiment (Fig 2).

Following the midway break, this process was repeated with the direction of a blue cursor misaligned with the direction of the participants’ unseen hand movement (Fig 1A+B). This misalignment was either a 30° or 60° CW visuomotor perturbation, depending on what group the participant belonged to. Participants were again, instructed to reach to targets as accurately and as quickly as possible. 90 trials of the reach training task were done at the start of the ‘Rotated’ session of the experiment and 30 trials were done during all other ‘Reach to Target’ tasks during the ‘Rotated’ session of the experiment (Fig 2).

The words “Reach to Target” were shown on the screen prior to the start of each set of ‘Reach to Target’ tasks to cue the participant of the next task.

#### Reach with No Cursor tasks

These tasks were completed at the end of each block. Participants reached to one of three targets much like in the ‘Reach to Target’ task. However, the cursor that indicated the position of the thumb was not visible and participants were asked to reach to the target without this visual feedback. When participants believed that they had acquired the target, they held their hand in place for 500 ms, indicating the completion of the reach, and the target disappeared. Participants then moved the robot handle back to the starting position along a robot-constrained path to begin the next trial. As in Werner et al. (2015), during ‘Rotated’ session of the experiment, during some trials, participants were asked to employ any strategy they used during the “Reach to Target” tasks. Only the participants aware of the rotation and how they were compensating for it were expected to show a clear distinction between reaches employing a strategy and reaches that do not since awareness of the perturbation is required to dissociate between the two conditions (Werner et al., 2015). Each of the three target locations was reached to three times during every ‘Reach with No Cursor’ task for a total of 9 reaches. The order in which ‘Reach with No Cursor’ tasks with and without strategy use were performed were counterbalanced within participants (between blocks, as shown in Fig 2) and between participants.

The words “No Cursor” were shown on the screen prior to the start of each set of ‘Reach with No Cursor’ tasks to cue the participant of the next task.

#### Localization tasks

Similar to the localization tasks used by Izawa et al. (2012), and ‘t Hart & Henriques (2016), participants were instructed to make outward movements while holding the robot handle with their unseen right hand in a direction of their choosing. The movements were stopped by a force cushion at a distance 10 cm from the starting position. Participants then moved the robot handle back to the starting position along a robot-constrained path. After moving back to the starting position, they were instructed to indicate the perceived location of their unseen hand at the end of their outward movement using a touchscreen located above the hand (Fig 1C+D). Importantly, they received no visual feedback of their right hand while their left hand, used only for localizing their right hand, was entirely visible.

For groups which trained to reach with a 30° rotated cursor during the ‘Rotated’ session, participants moved their unseen hand toward an arc located 10 cm away from the starting position that spanned 60 degrees (similar to the white arc in Fig 1D), centered on the 50°, 95° or 130° mark in polar coordinates. During each set of localization trials, the arc appeared six times in each of the three possible locations to encourage subjects to move their hand to a large range of locations. The participants indicated their perceived hand location by touching on these same arcs with their visible left hand. For groups which trained with a 60° rotated cursor, participants were instructed to move their unseen hand in a direction of their choosing that fell within a visible V shaped wedge (Fig 1D), with the tip of the wedge at the starting position and the two ‘arms’ pointing outward. This wedge was again used to encourage participants to move to a large range of locations without providing visible targets that may bias touch-screen responses. After completing the movement, an arc appeared 10 cm from the starting position that spanned from 0° to 180° in polar coordinates and participants indicated their felt hand position by touching where they perceived their unseen hand to have crossed that arc. Participants placed their left hand under their chin between taps on the touch screen to avoid confounding contact with the touch screen.

In all ‘Active Localization’ tasks, participants made volitional movements to a location of their choice. After the robot handle was moved 10 cm from the starting position, a force was applied to prevent the participant from reaching further, giving them the sensation of hitting a wall. In passive versions of the localization task, adapted from the task in ‘t Hart et al. (2016), the participants’ unseen right hands were pulled by the robot manipulandum to various points on the arc. These points on the arc were either identical to the points to which the participants actively reached in the preceding active version of this task or located at the center point of the arcs (there was no significant difference in localization in either case). Like the active localization task, after hitting a point on the arc, the participants were instructed to return their unseen right hand to the starting position and then indicate with their visible left hand on a touch screen where the right hand intersected with the arc. We find no effects of the positions on the arc which participants’ hands were moved to during the passive version of task on any of our primary measures. Therefore, to better analyze changes in hand-localization, we collapse across these conditions and treat them as one group.

The words “Right Hand: Cross Arc, Left Hand: Touch Cross” or “Robot: Cross Arc, Left Hand: Touch Cross” were shown on the screen prior to the start of each set of localization tasks to remind the participant to indicate the part of the arc which they thought they crossed during either self-generated or robot-generated movements.

#### Questionnaire

After completing the experiment, participants were asked a series of questions (Fig 3) similar to that of Benson et al (2011). The questions were used to determine whether participants were conscious of the perturbation and to assess how accurately participants perceived they compensated by using the explicit strategy.

**Fig 3:**
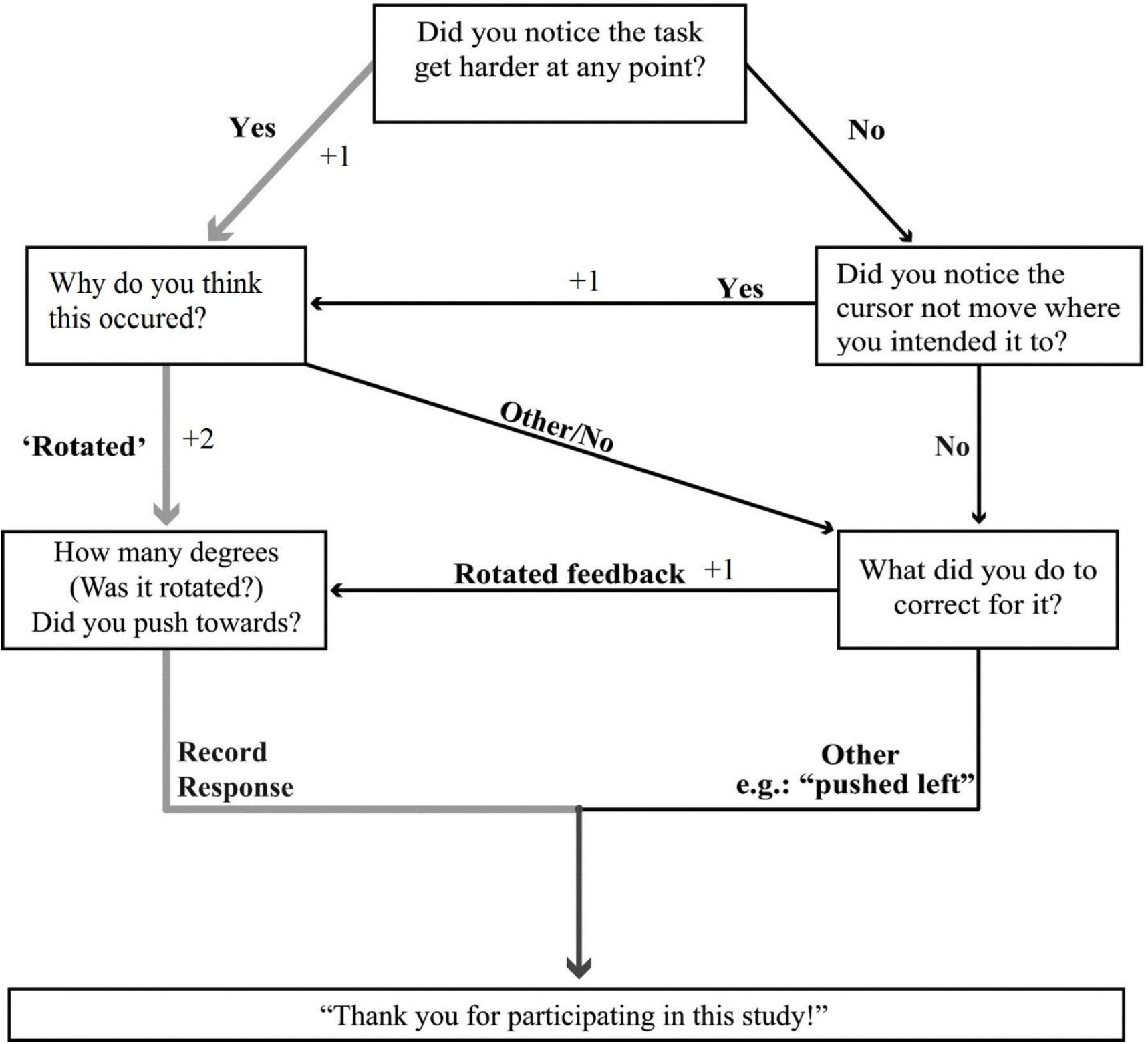
Post-experiment questionnaire. These questions were asked regarding the ‘Reach to Target’ tasks. Follow-up questions were determined by answers to the previous question. Each answer was given a score to attain an ‘Awareness Rating’ for each participant, with a maximum rating of 3 and a minimum rating of 0.

### Analysis

We aimed to determine the effects of awareness of a visuomotor perturbation during reaching movements on adaptation-related changes in hand localization. To ensure that participants who were instructed or experienced large rotations were in fact aware and used a strategy when adapting to the perturbation, we analyzed reaching movements both during ‘Reach to Target’ tasks and ‘Reach with No Cursor’ tasks. During each reaching movement, both with and without a visible cursor-representation of the hand, angular hand deviations were calculated at the point of maximum hand velocity. The angular deviation was the angular difference between a line from the starting position to the target and a line intersecting both the starting position and the position of a participant’s hand at the point of maximum velocity. When reaching with a rotated cursor representation of the hand, participants had to deviate their reaches by either +30° (NI30 and I30 groups) or +60° (NI60 and I60 groups) to fully compensate for the rotation, where positive refers to CCW direction. To make performance during training comparable across the two rotation groups, we also normalize the hand deviations to the size of the experienced rotation.

Estimates of hand location were determined by the angular difference between the end-point of the unseen right-hand movement, and participants’ perceived hand position as indicated on the touchscreen. We used a kernel smoothing method to represent hand-localization changes across the workspace. We used a normal kernel with a width of 10 degrees to interpolate the shifts in localization at specific hand-movement angles; 50, 90 and 130 degrees in polar coordinates. These angles were the center points of guiding arcs and wedges in the ‘Localization’ tasks. We then used the means of these interpolated values for all statistical analyses involving hand-localization changes. Given that the cursor was rotated CW, localization shifts should also be in the CW, or negative, direction.

First, we analyzed the effects of instruction and rotation size on the process of learning during the first 90 trials of rotated-cursor training. We subtracted individual biases in reaches during the last 45 trials in the aligned session from the hand deviations recorded in the rotated session, and blocked the training data into trial sets of 3 trials each for the first 6 trials of rotated-training, and a final trial set of the last 9 trials of training. We then performed a 2×2×3 mixed ANOVA with instruction (*instructed* or *non-instructed*) and rotation size (*30°* or *60°*) as between-subject factors, and trial set (*trial sets 1, 2, or final* during training) as a within-subject factor to examine the effects of instruction and rotation size on performance changes during adaptation. To examine this more closely, we performed 2×2 ANOVAs with instruction and rotation as factors on each of the three analyzed trial sets. All tests had an alpha level of 0.05 and applied Greenhouse Geisser corrections where necessary.

Next, we assessed the use of explicit strategies during adaptation using the process dissociation procedure (PDP), where participants reached to targets without cursor-feedback. The PDP was adapted from a study by Werner et al. (2015) and was used to measure awareness of the perturbation after adaptation. In the PDP, we calculated mean hand deviations per each participant when performing reaches while employing any strategy they used during adaptation and when not employing any strategy. First, we determined if adapting to a visuomotor rotation led to changes in hand deviation, that is, reach aftereffect, in ‘Reach with No Cursor’ tasks, for those trials where they were told not to employ a strategy. We ran a 2×2×2 ANOVA on these no cursor reaches with session as a within-subject factor, and instruction and rotation size as between-subject factors to confirm that training did lead to reach aftereffects, and to further test if these reach aftereffects changes with instruction or rotation size. After confirming that implicit motor changes (reach aftereffects) occur after adaptation, we subtracted individual biases in hand deviation in the ‘Reach with No Cursor’ during the aligned session from those in the rotated session. Using these baseline-corrected hand deviations, we ran a 2×2×2 ANOVA on PDP reaches (‘Reach with No Cursor’ tasks during the rotated session) with instruction and rotation size as between-subject factors, and strategy use as a within-subject factor to examine the effects of instruction and rotation size on performance in the PDP. Awareness of the perturbation would be associated with a significant difference between the two types of Reach with No Cursor, while lack of awareness would be associated with no difference.

To quantify this awareness of perturbation in this PDP across our groups, we calculated a ‘Strategy Use Ratio’ for each participant using their hand deviations in the PDP tasks:

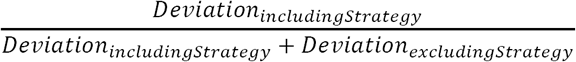

Participants who were not aware of the perturbation, and thus could not employ a strategy when asked, were expected to have a Strategy-Use Ratio near 0.5. Participants who were more aware, and who used an explicit strategy during adaptation were expected to have higher Strategy-Use Ratios. Participants whose adaptation was entirely explicit, meaning they could entirely suppress hand-deviation when asked to exclude any strategies, were expected to have a Strategy-Use Ratio of 1.0.

To answer our question whether awareness affects changes in hand localization, we analyzed how afferent and efferent based changes in hand localization due to adaptation were independently affected by instruction and rotation size. First, we wanted to confirm that hand localization shifted with adaptation to a visuomotor rotation, and whether volitional movements in the ‘Active Localization’ tasks lead to additional localization changes. To do so, we compared participants’ estimates of hand position before and after adapting to the perturbations using a 2×2 ANOVA with session (*aligned* or *rotated*) and movement type (*active* or *passive*) as factors. After confirming hand localization changes due to adaptation, we subtracted individual localization biases in the aligned session from those in the rotated session. We then isolated afferent based changes in hand localization (changes measured in ‘Passive Localization’ tasks where participants did not have access to efferent signals) and efferent based changes in hand localization (the difference between changes in ‘Active Localization’ tasks, where participants did have access to efferent signals, and ‘Passive Localization tasks). This was followed by two 2×2 ANOVAs with instruction and rotation size as between-subject factors on both sets of data.

Finally, we explored the relationships between awareness of the perturbation, changes in movements, and hand estimates,, by comparing measures in the ‘Reach with No Cursor’ tasks, i.e. the PDP, with those in the localization tasks. To assess relationships between either afferent or efferent-based shifts in hand localization and the Strategy Use Ratios of individuals, we computed Pearson product-moment correlations. Likewise, we also computed Spearman’s correlations to determine relationships between either afferent or efferent-based shifts in hand localization and Awareness Scores on post-experiment questionnaires. We computed additional Pearson product-moment correlations to determine relationships between either afferent or efferent based changes in hand localization and hand deviations in ‘Reach with No Cursor’ tasks with and without strategy use.

Data preprocessing was done in Python version 3.6 and data analysis conducted in R version 3.4.4.

## Results

### MANIPULATING AND MEASURING AWARENESS OF THE PERTURBATION DURING ADAPTATION

When exposed to a visuomotor rotation, participants in all groups learned to deviate their reaches to counter ∼87% of the rotation by the end of 90 training trials. To examine the effects of instruction and perturbation size on the extent of our participants’ adaptation, we analyze the first trial set of training where we expect to see the largest effects. When hand deviations were normalized relative to the rotation size (Fig 4B+C), we find that only instruction (main effect; F(1,80) = 81.245, p < .001, generalized eta squared (η^2^_*G*_) = .504), and not the rotation size (main effect; F_(1,80)_ = 0.031, p = .861, η^2^_*G*_ < .001) lead to greater initial compensation. As illustrated in figure 4B, instructed groups adapted similarly and non-instructed groups adapted similarly over the 90 training trials. The difference in hand-deviation due to instruction persisted throughout the second trial set of training (main effect; F_(1,80)_ = 15.837, p < .001, η^2^_*G*_ = .165) and even until the end of the training task (main effect; F_(1,80)_ = 5.781, p = .019, η^2^_*G*_ = .067). In short, as seen in Fig 4, when instruction but not rotation size led to more substantial compensation during training.

**Fig 4:**
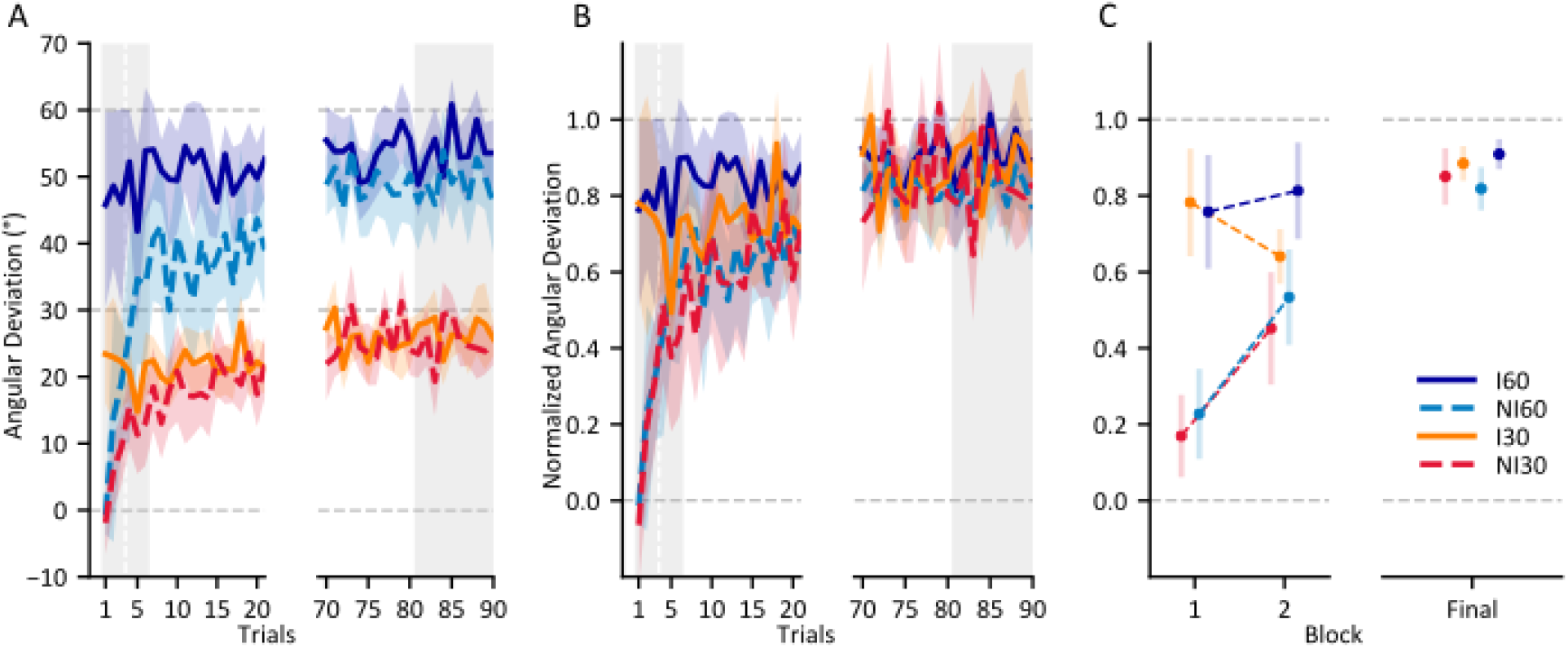
Performance of all groups during adaptation. A) Hand deviations in the direction countering the perturbation for the first and last 20 trials of training. The indicator lines at 30 and 60 degrees demonstrates the hand deviation required to fully counter the perturbation for groups that experience a 30 and 60-degree rotation respectively. B) Results in A normalized with respect to rotation size. The indicator line at 1.0 demonstrates full compensation of the perturbation. C) Normalized mean hand-deviations for the first two three-trial sets of training and the final nine trials of training used in the ANOVAs, as indicated by the grey area in A and B. Shaded areas and error bars are 95% confidence intervals.

Next, we confirmed that adapting to a rotated cursor lead to significant reach aftereffects (continued hand deviations when reaching without a visual cursor representation of the hand). That is, hand-deviations in the ‘Reach with No Cursor’ tasks where participants were not told to use an explicit strategy were deviated 14.1° on average in the direction countering the visuomotor rotation in the rotated session when compared to the aligned session (main effect of session; F_(1,80)_ = 509.295, p < .001, η^2^_*G*_ = .680). These implicit aftereffects of adaptation were equal in size for all four groups (Fig 5A: Without Strategy), independent of instruction and even the size of the rotation (no interactions between session and either instruction or rotation size). This suggests that implicit aftereffects are not suppressible and are rigid in their magnitude, regardless of differences in the size of the experienced perturbation, and even despite awareness of perturbations during adaptation.

**Fig 5:**
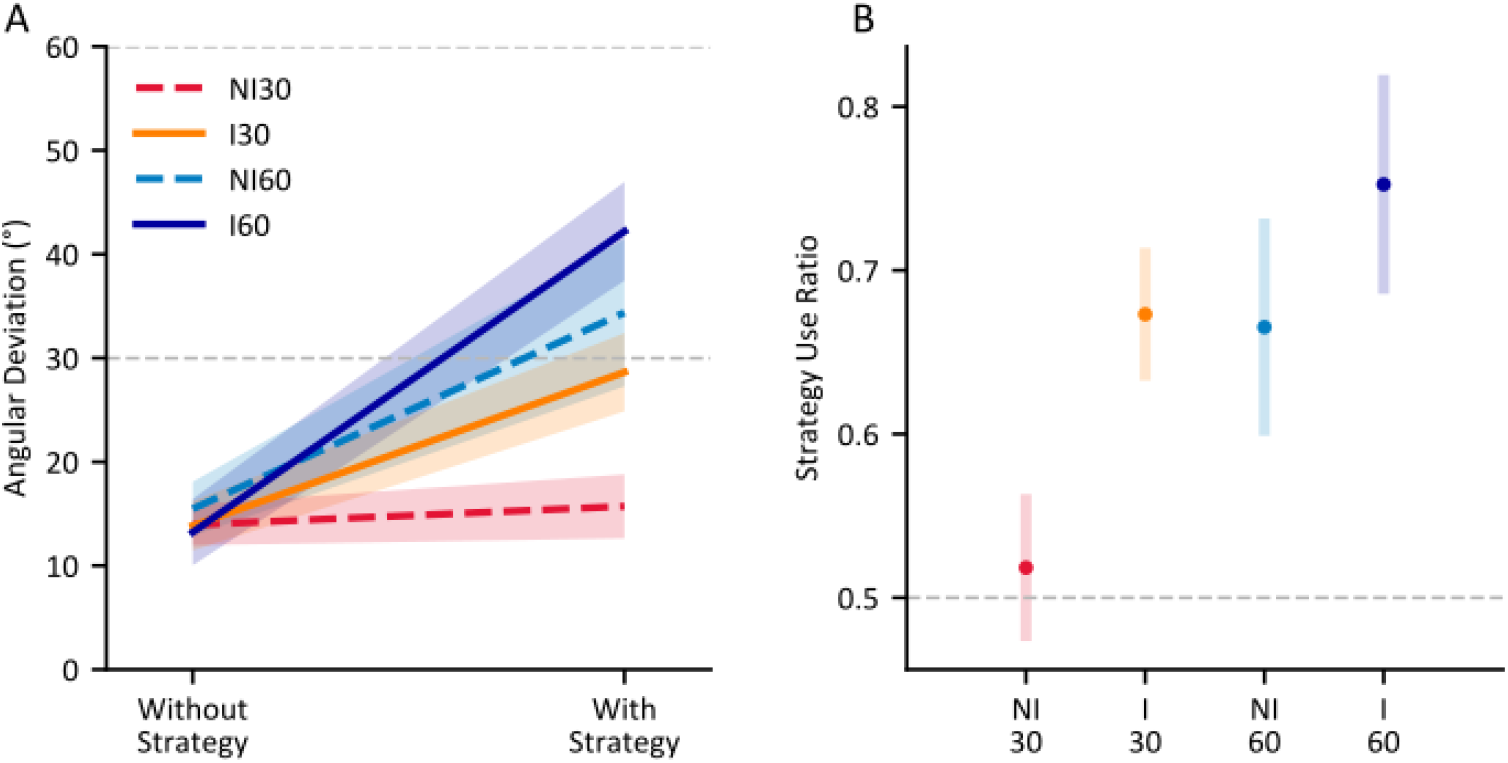
Changes in no-cursor reaches following training. A) Mean group deviations in movement direction, while suppressing (left side) or using (right side) any strategies employed during adaptation. B) Mean Strategy Use Ratios per group, calculated using differences in movement direction when employing and not employing a strategy. 0.5 indicates an inability to differentiate between the two conditions; this should reflect awareness. Shaded areas and error bars are 95% confidence intervals.

To test our prediction that both strategies and large perturbation sizes would result in awareness of the perturbation during adaptation, we examine the ‘Reach with No Cursor’ trials where participants use explicit strategies they employed during adaptation (Fig 5A: “With Strategy”) and compare them with trials where participants do not use any strategy. We tested whether participants were able to evoke a strategy when cued during the PDP, i.e., to further deviate their hand in reaches with strategy use when compared to reaches without strategy use. Reach deviations with a strategy were larger than those produced when asked to exclude any conscious strategy, but only for some of the groups. Specifically, being instructed (instruction ^*^ strategy use interaction; F_(1, 80)_ = 17.131, p < .001, η^2^_*G*_ = .106) and experiencing a large perturbation (size ^*^ strategy use interaction; F_(1, 80)_ = 31.476, p < .001, η^2^_*G*_ = .179) led to greater hand deviations when asked to use strategy than when asked not to. This suggest that training with a larger rotation even without instruction is sufficient to develop awareness of the nature of the perturbation. As illustrated in Fig 5B, only the NI30 group (red) did not show significant strategy use (Strategy Use Ratios; M = 0.518, SD = 0.096, one-sample t-test against 0.5: t(19) = 0.855, p = .202). However, although participants who experience a large perturbation developed strategies to counter it, neither group which adapted to a 60° rotation could employ a strategy to counter the full magnitude of the rotation (NI60 countered 70% and I60 countered 77% of the rotation when asked to use strategy). Our results suggest that either receiving instruction or training with a large perturbation can lead to awareness of the perturbation, as well as a strategy for how to counter it, during adaptation to visuomotor rotations, but there may be other factors at play when countering large perturbations.

### THE EFFECTS OF AWARENESS DURING ADAPTATION ON HAND-LOCALIZATION

Having confirmed that instruction and large perturbation sizes lead to increased awareness of the nature of the perturbation during adaptation, we examine the effects of differences in awareness on changes in estimates of hand position. Specifically, we probe both afferent (via available sensory information) and efferent (via an efference copy of produced motor commands) based changes in localization of the hand following adaptation. To isolate efferent and afferent based changes, we use one localization task in which participants have access to both efferent and afferent based signals of hand location (the ‘Active Localization’ task) and another in which participants only have access to an afferent based signals of hand location (the ‘Passive Localization’ task). By comparing the two tasks, we attempt to probe changes in hand localization based only on efferent based information.

All groups showed a change in hand localization following adaptation (main effect of session (aligned vs rotated); F_(1, 83)_ = 172.285, p < .001, η^2^_*G*_ = .171). These changes were modulated by the type of movement of the hand being localized (session * movement type interaction; F_(1, 83)_ = 28.374, p < .001, η^2^_*G*_ = .005), suggesting a further shift in efferent-based perceived hand position when the hand was actively moved by the participant. A majority of the observed shift, 5.5°, was afferent-based, present in the ‘Passive Localization’ task. Neither instruction nor rotation size affected this proprioceptive recalibration of hand localization (F_(1, 80)_ = 0.254, p = .616, η^2^_*G*_ = .003) and F_(1, 80)_ = 2.295, p = .134, η^2^_*G*_ = .028 respectively). That is, proprioceptive recalibration was not modulated by instruction, and saturated at around 5.5°, no matter how large the perturbation or adaptation.

When we isolate efferent-based shifts in hand localization, we find 2.1° of additional shifts (about 38% larger) to the afferent based changes present in ‘Passive Localization’ tasks (Fig 6: panel A vs B, illustrated in panel C). These additional shifts can be attributed to updating of efferent based estimates of hand position. However, as for afferent based changes, neither instruction (F_(1, 80)_ = 3.958, p= .050, η^2^_*G*_ = .048) nor rotation size (F_(1, 80)_ = 0.007, p = .936, η^2^_*G*_ < .001) significantly modulated these changes. Thus, our results show that neither instruction nor rotation size have measurable effects on changes in either efferent or afferent based hand localization (Fig 6).

**Fig 6:**
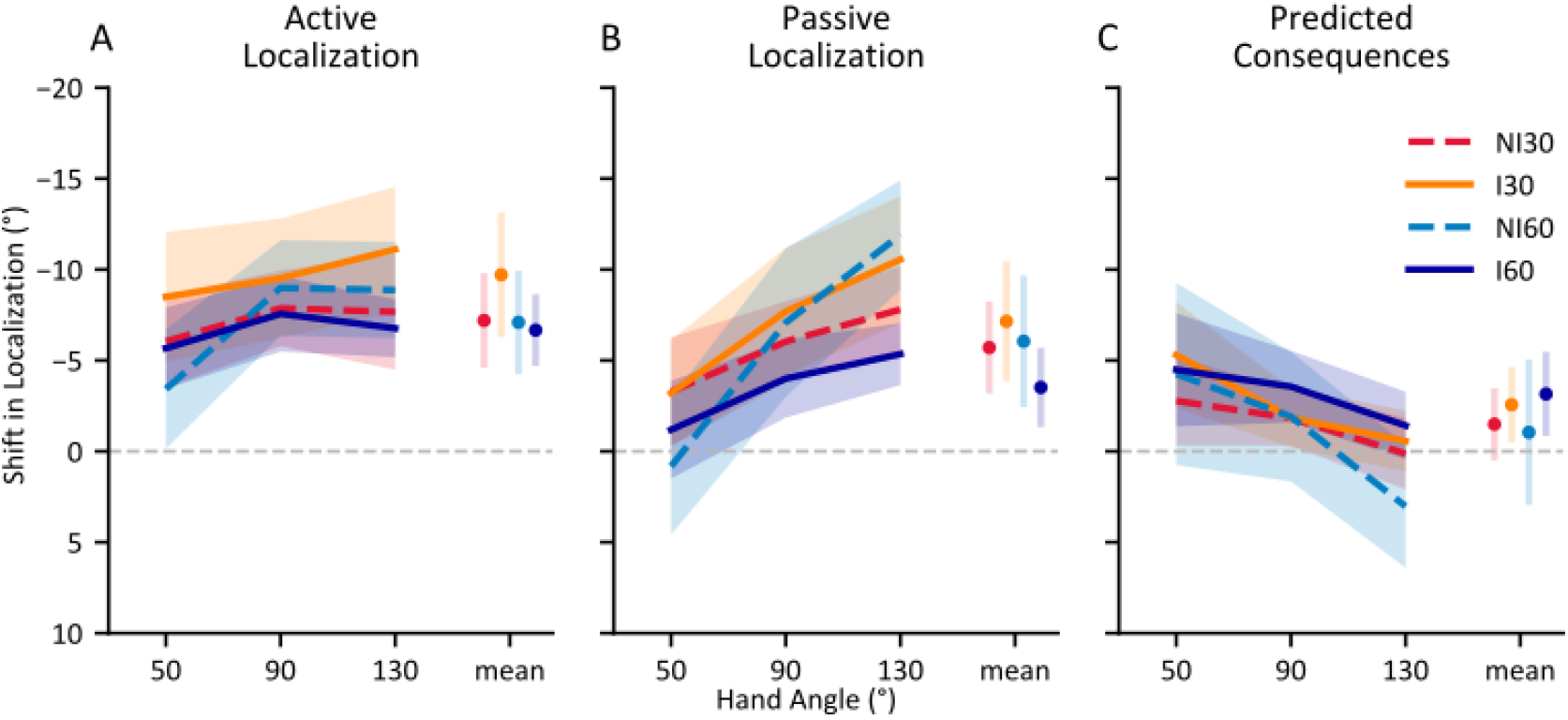
Changes in localization of the unseen, adapted hand following visuomotor adaptation. A) After self-generated movements; Active Localization B) After robot-generated movements; Passive Localization C) Difference between Active and Passive Localization, intended to capture updates in predicted sensory consequences (efferent based estimates). Shaded areas and error bars are 95% confidence intervals.

Self-reported questionnaire responses were poor representations of awareness levels when compared to behavioural measures such as the PDP (Werner et al. 2015). We added scores to our questionnaires to improve the efficacy of self-reported questionnaires. When assigning a 4-point awareness score to the responses on the questionnaire used in Benson et al. (2011), (see Fig 7A) we find that higher scores on the questionnaire positively correlate with an individual’s Strategy Use Ratio (r_s_ *=*0 .373, p *<* .001), but this is partially due the clustering of certain groups in both the awareness scores and Strategy Use Ratios.

**Fig 7:**
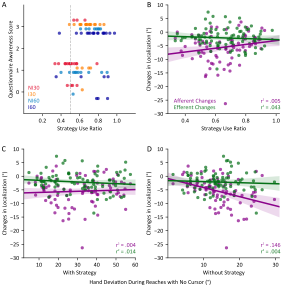
Relationships between Strategy Use Ratios, Questionnaire Awareness Scores, and changes in hand localization. (A) Individual participants’ questionnaire awareness scores compared to their Strategy Use Ratios. (B) Relationships between afferent and efferent based changes in hand localization and Strategy Use Ratio. (C+ D) Relationships between afferent and efferent based changes in hand localization and reach deviations without a cursor representation of the reaching hand while employing(C) and not employing (D) any explicit strategies used during adaptation.

To ensure that awareness of the perturbation during adaptation does not effect changes in felt hand position at an individual level, we also test whether a participant’s Strategy Use Ratio or Awareness Scores are correlated with their changes in hand localization. Consistent with our analysis above, Strategy Use Ratios were not correlated with either afferent based (r = .207, p = .059) nor efferent based (r = -.069, p = .530) changes in hand-localization (Fig 7B). Likewise, awareness scores from self-reported questionnaires (Fig 3) also did not correlate with either afferent based (r_s_ *=* .092, p *=* .404) nor efferent based (rs = -.063, p *=* .567) changes in hand-localization (not shown). Furthermore, we find that afferent and efferent-based changes in hand localization were also not correlated with participants’ hand-deviations during ‘Reach with No Cursor’ trials in which they use an explicit strategy (Fig 7C: r = .059, p = .059 and r = -.120, p = .278 respectively). However, as illustrated in Fig 7D, afferent based localization changes were significantly correlated with hand-deviations during ‘Reach with No Cursor’ trials where participants did not use an explicit strategy, i.e., implicit motor changes or reach aftereffects due to adaptation (r = -.382, p < .001), although efferent-based localization changes were not (r = -.061, p = .580). Our findings suggest that, for the most part, efferent and afferent changes in hand localization, and motor changes due to adaptation are separate processes, but there may be similar mechanisms between afferent-based changes in localization and implicit motor aftereffects of adaptation. That is, proprioceptive recalibration partly predicts implicit motor changes.

Our measures of awareness show we were successful at evoking explicit awareness of a perturbation during adaptation, both by instructing participants and by having them adapt to large perturbations. Our results taken together indicate that changes in hand-localization due to adaptation, both when relying on afferent-based and efferent-based information are largely implicit processes. High awareness levels in various awareness measures, i.e., participant questionnaires and process dissociation procedures, do not result in any significant changes in the shifts in localization of the adapted hand. That is, implicit components of motor adaptation, including the localization changes, are rigid, found in all groups, independent of cognitive factors.

## Discussion

We tested whether awareness of a perturbation during adaptation, brought about by either instruction or by experiencing a large 60° visuomotor rotation, can modulate changes in the estimation of hand position. We find that adapting to a visuomotor rotation lead to significant changes in hand localization, which are both afferent based, informed by sensory information from the effector, and efferent based, informed by a copy of a motor command during a movement. We find that both instructions and large rotations affect measures of explicit learning, but they do not impact changes in estimates of hand position. As we will discuss below, our findings have implications for the processes involved in both proprioceptive recalibration and updating of predicted sensory consequences.

### EXPLICIT LEARNING AND AWARENESS OF THE PERTURBATION

Contributions of explicit and implicit components to motor learning have been measured both directly, by having participants indicate their employed aiming strategies when reaching with rotated visual feedback (Bond & Taylor, 2015; Hegele & Heuer, 2010; Heuer & Hegele, 2008; Heuer & Rapp, 2011; Taylor, Krakauer, & Ivry, 2014), and indirectly, using aftereffects of reach adaptation or post-experiment questionnaires for instructed and non-instructed groups (Benson et al., 2011; Neville & Cressman, 2018; Werner et al., 2015). Where measured directly, explicit learning appears to dominate in early session s of adaptation, when errors are large and salient (Bond & Taylor, 2015; McDougle et al., 2015, 2016; Taylor et al., 2014). Consistent with these previous findings, although we find an explicit component of learning due to instruction that is present up to 90 trials into training to counter a visuomotor perturbation, we see the largest benefits of instruction during early learning, within the first 6 trials. The size of the effect of instruction during late adaptation was smaller (η^2^_*G*_ = 0.067) when compared to the first 6 trials of rotated training (η^2^_*G*_ = 0.503, and η^2^_*G*_ = 0.165 for the first and second trial sets respectively). Participants who experienced a larger perturbation but did not receive instructions adapt at a faster absolute rate, but these changes in hand deviations are proportional to the rotation size. As adaptation progresses, the contribution of explicit learning continues to decrease as implicit aspects of learning slowly increase and begin to dominate the adaptation process (Taylor et al., 2014). Although we do not directly measure explicit components of learning during training, like Benson et al (2011), Werner et al. (2015) and Neville & Cressman (2018), we see evidence for potential benefits of having well-informed cognitive strategies during early adaptation. That is, our results suggest instruction on the nature of a perturbation may lead to increased explicit components of learning even when perturbations are not salient.

Although performance as a whole is similar between people who experience the two rotation sizes when normalized, measures of explicit components of adaptation, including awareness of the perturbation following adaptation, have been repeatedly shown to differ with the size of the perturbation (Bond & Taylor, 2015; Hegele & Heuer, 2013; Heuer & Hegele, 2008; Neville & Cressman, 2018; Werner et al., 2015). We measured the consequences of these explicit components of learning indirectly following reach training, using the process dissociation procedure (PDP: Fig 5) and found that people become aware of a 60° visuomotor perturbation and develop a strategy to counter it when adapting to it naïvely. A 30° rotation on the other hand, was not sufficient in evoking the use of a conscious strategy throughout our 90 trials of training. As with our participants, people could evoke a strategy when cued after experiencing large perturbations of 75° (Hegele & Heuer, 2013), 60° (Werner et al. 2015; Neville & Cressman, 2018) and 40° (Neville & Cressman, 2018). In contrast, people could not consistently evoke a strategy after adapting to perturbations of 20° (Werner et al. 2015; Neville & Cressman, 2018) or 40° (Werner et al. 2015). The factors effecting the lower bounds of the rotation size required to develop lasting awareness of the perturbation are still unknown. Conversely, when explicit components of learning were directly solicited during training, even rotation sizes as small as 15° were sufficient for the development of a strategy which persisted throughout training (Bond & Taylor, 2015). Although direct measures of explicit learning are informative about its expression during adaptation, the task of choosing an aiming direction prior to reaching may lead to an overexpression of explicit learning (Werner et al., 2015). By using the PDP, we avoided overexpression of explicit components in non-instructed groups. Using this method, we show that non-instructed 30° participants were not aware of the perturbation whereas those that experience a 60° perturbation were.

A reliable method of evoking awareness of the perturbation during training is simply informing the participants of the perturbation and how to counter it. When we provided such instruction to participants, we observed increased strategy use in people that adapted to both large and small rotations (Fig 5B: I30 and I60 groups). Indeed, for a small rotation of 30°, instructed participants on average evoke a complete strategy, showing hand deviations that matched the perturbation, in the absence of the cursor. For the larger 60° rotation, hand deviations produced when evoking the strategy were not nearly as complete, although this is generally the case seen in other studies (e.g.,(Hegele & Heuer, 2013). Like in our study, Werner et al. (2015) and Neville & Cressman (2018) found that providing instruction increased the ability to evoke an explicit strategy when cued during a PDP task, regardless of the size of the experienced perturbation. As with these previous studies, our PDP measures show that instructing participants and having them experience large perturbation sizes each lead to awareness of the perturbation, with slightly greater hand deviations for the group that had both (I60); although this does not reach statistical reliability. Overall, by using post-training measures of perturbation awareness, we were successful in evoking awareness in some groups (I30, NI60, I60) and not others (NI30), allowing us to cleanly study its effects on other consequences of motor adaptation.

Implicit components of adaptation are clearly independent of explicit components, differing both in the time course of their development during training (McDougle et al., 2015; Taylor et al., 2014) and in their development during the aging process (Heuer & Hegele, 2008). They are however related, as patients with cerebellar ataxia show deficits in both implicit and explicit aspects of reach adaptation (Butcher et al., 2017). We find that reach-aftereffects made without strategy were consistent regardless of instruction or the size of the perturbation. As in our study, Bond and Taylor (2015) found that implicit aftereffects were consistent among groups after adapting to a single rotation size. However, others have shown that implicit aftereffects can be proportional to the rotation size, but these cases tended to either gradually introduce the perturbation (Salomonczyk et al., 2011; Salomonczyk, Cressman, & Henriques, 2013) or include multiples rotation sizes during training (Thomas & Bock, 2012; Werner et al., 2015). Introducing multiple rotation sizes or introducing a perturbation gradually may increase implicit components of learning by exposing the participant to smaller required adjustments. We can only speculate at the mechanisms as this is beyond the scope of this study. While we find no effects of instruction on implicit reach aftereffects, Neville & Cressman (2018) found that providing instruction lead to smaller implicit after-effects of learning, measured by the PDP. These discrepancies may be related to the specifics of instructions given to participants both before adaptation, and before performing the PDP task.

### HAND LOCALIZATION

The roles of instruction, strategies and explicit learning on resulting changes in hand localization have not been explored. While it has been shown that motor adaptation leads to both afferent and efferent based changes in hand-localization (’t Hart & Henriques, 2016; Cressman & Henriques, 2009), it is not clear if the changes are entirely implicit, or if they can be modulated by explicit components of learning. Awareness of the perturbation may lead to experienced errors being assigned to an external source rather than one that is internal, which has been shown to affect adaptation (Berniker & Kording, 2008; Kong, Zhou, Wang, Kording, & Wei, 2017; Wilke, Synofzik, & Lindner, 2013), and thus should not lead to changes in body-based estimates. However, regardless of whether participants are given instructions prior to adaptation, and regardless of whether they exhibit awareness of the perturbation following training, participants did not show any effects of instruction or rotation size on changes in either proprioception or efferent-based hand localization.

Moreover, the size of these shifts in localization did not vary with the size of rotation. This is in contrast to an earlier study from our lab (Salomonczyk et al., 2011). Gradually introducing the cursor rotation and subsequent increases in its rotation did lead to proportionally larger changes in hand estimates (as well as implicit reach aftereffects as mentioned above) after sufficiently training with each of the three final rotations; roughly accounting for 20% of rotations of 30, 50 and 70 degrees. Gradually introducing the different rotations may have made the perturbation less “explicit” to participants in that study, therefore resulting in larger changes to both proprioception and reach aftereffects. Thus, the abrupt nature of the 60° perturbation in our study may have limited implicit proprioceptive and motor changes to a more tolerable level, similar in magnitude to those produced during implicit learning of the smaller rotation. Recently, implicit motor effects of learning have been shown to be insuppressible, developing even when participants have no control over the direction of cursor movement in a reaching task and are instructed ignore the cursor error (Kim, Morehead, Parvin, Moazzezi, & Ivry, 2018; Morehead, Taylor, Parvin, & Ivry, 2017). Our study suggests that, like the motor aftereffects of adaptation, changes in perceived hand location are also insuppressible and may rely on similar, but distinct mechanisms. Specifically, changes in afferent-based hand localization, i.e., proprioceptive recalibration, have been shown to be separate from motor changes; with different time courses (Ruttle et al., 2016) and generalization patterns (Cressman & Henriques, 2015). Since changes in localization are not affected by our manipulations of awareness of the perturbation, they might be largely implicit processes. The modest relationship between implicit motor and proprioceptive recalibration (Fig 7D), further supports that both are similarly unconscious processes.

Hand localization, and their shifts due to motor adaptation are informed both by afferent signals, such as proprioception, and efferent signals, like the motor commands to generate volitional movements, the latter of which may be altered by the changed internal forward model after adaptation (Mazzoni & Krakauer, 2006; Tseng et al., 2007). Synofzik et al. (2008) and Izawa et al. (2012) have found significant changes in estimates of hand position, using an active localization task, following adaptation (Izawa, Criscimagna-Hemminger, & Shadmehr, 2012; Shadmehr, Smith, & Krakauer, 2010; Synofzik, Lindner, & Thier, 2008). As replicated in our study, when a passive version of the localization task was used to isolate afferent-based changes and efferent-based changes in hand localization, around 80% of a 10° shift in hand localization could be attributed to changes in afferent-based localization, with only 20% that can be attributed to efferent based signals (’t Hart & Henriques, 2016). This is consistent with the additional results of Synofzik et al. (2008) and Izawa et al. (2012), where they found that the learning-induced shifts in active hand localization was reduced (although still significant) in patients with cerebellar damages, and our own results that show cerebellar patients do show proprioceptive recalibration (Henriques, Filippopulos, Straube, & Eggert, 2014). This suggest that indeed the cerebellum may be contributing to predicting the sensory consequences of movements but not proprioceptive estimates of hand position (Block & Bastian, 2012; Henriques et al., 2014). We also isolated these components and observed how they interact with awareness, and indirectly source attribution of errors, during adaptation. When we isolated these efferent based estimates of hand position, they, much like afferent based changes, were also not modulated by awareness of the perturbation. Our finding suggest that these efferent based changes too are implicit in nature.

### MEASURING AWARENESS

When determining awareness, relying solely on questionnaires may lead to its underestimation as verbal and motor responses are significantly different retrieval contexts (Shanks & St John, 1994). Furthermore, questionnaire responses on higher order cognitive processes are held in low confidence, as responses may be effected by multiple factors including the saliency of stimuli related to the response, levels of attempted introspection and even conscious access to cognitive components of prior performance (Eriksen, 1960; Nisbett & Wilson, 1977). Thus, we used a variation of a process dissociation procedure (PDP), adapted by Werner et al. (2015) to objectively measure awareness following adaptation. Werner et al. (2015) found that PDP results were informative, related to performance during adaptation and during catch trials which measured implicit adaptation, whereas binary questionnaire results were not. Although the PDP may be a more principled method of measuring awareness, we find that adding a scoring system to the questionnaire (Fig 3) used by Benson et al (2011), where different degrees of awareness are accounted for, can provide insights on participant awareness (Fig 7A). Participants’ scores on the modified questionnaire correlate to the PDP measures, while no such relationship arises when the outcome of the questionnaire is dichotomous (Werner et al., 2015). In an ambiguous situation, as in the NI60 group where some participants develop an explicit strategy, questionnaire responses, although non-tractable, may be an asset when performance in the PDP is not measurable.

## CONCLUSION

Having instructions on the nature of a visuomotor perturbation before experiencing it, as well as experiencing a large perturbation, lead to the development or use of an explicit strategy during adaptation. Even when adaptation involves these explicit processes, implicit aftereffects of adaptation, in the form of continued deviated reaches in the absence of a perturbation, continue to be present. The magnitude of these aftereffects is independent of the size of the perturbation, and whether participants are given a strategy to counter the perturbation. Adapting to a visuomotor rotation leads to changes in estimates of unseen hand position. Estimates, which are based both on afferent information, from sensory inputs such as proprioception, and efferent information, derived from efference copies of motor commands, both significantly shift to more align with visual input during adaptation. These changes in hand localization are not modulated by awareness of a perturbation or the use of a strategy during adaptation. These findings support the notion that both proprioceptive recalibration and efferent based changes in hand-localization are insuppressible and largely implicit in nature, and like other implicit components of motor learning, develop independently of explicit components, and possibly from each other.

## APPENDIX

### Appendix A: Instructions received by participants prior to and throughout the experiment

#### Intro

In this experiment you will be completing a variety of tasks while using your right hand to grip the handle of a robotic arm (place thumb on top of screw). The robot handle can be moved by you or it can mechanically guide your movements, depending on the task. At some points you will be responding to stimuli displayed on the touch screen in front you and using your left hand to touch certain spots reached by your right hand.

#### Cursor (“Reach to Target”)

The green <u>cursor</u> on the screen represents your hand position – specifically the thumb. You are going to be reaching to the yellow <u>target</u> while gripping and moving the robot handle. The cursor has to overlap with the target for the trial to be complete. After completing the reach, you wait for the target to disappear and then move the handle back to the starting position. Then you can remain still and wait for the next trial to begin. Try to move your hand to the target as quickly and accurately as possible.

#### Active Trials (“Right Hand: Cross Arc…etc.”)

In this task you are going to see an arc. Your task is to reach with your unseen right hand (while gripping the robot handle) to intersect the arc at a point you choose. As this task repeats try to reach to intersect different sections of the arc instead of just one point. The more areas of the arc you intersect the less time this experiment will take. After intersecting the arc, you pull your arm back to the starting position. Next, you will use your left hand to indicate on the touch screen where your right hand crossed the arc. Once you have touched on the screen with your left hand, retract your hand and place it under your chin (demonstrate).

#### Passive trials

In this task you are going to see an arc just like before. However, the robot will move your unseen hand to a specific part of the arc. Let the robot guide your arm movement; don’t resist. As before, you will indicate where your right hand crossed the arc, with your left hand and remove the left hand from the touch screen in between trials.

#### No Cursor

In this task you are going to reach to targets but this time there is no cursor indicating your hand position. Your task, like before, will be to reach to the target as accurately as possible. After completing the reach – when you think you have reached the target – keep your hand still. Keeping your hand still informs the robot the trial is complete, and the target will disappear. Then move the handle back to the starting position and wait for the next trial to begin.

#### For EXPLICIT version only: Explaining Perturbations/Rotation task

For these next few reaching tasks, the green cursor will move a bit differently. The cursor is not going to move in the same direction as your hand and you will need to compensate for this. [SHOW and explain DEMO]. Imagine your starting point as being at the centre of a clock face, and you move your hand to the 12 position at the top. As you move your hand from the centre to the 12, the cursor will move to the right to the 2 on the clock. Aim for the 10 on the clock so that your cursor ultimately reaches the target at 12. Did I explain this clearly?

Keep these instructions and strategy in mind since you will be asked to use this strategy several times, including when reaching without a cursor. Sometimes you will be asked to NOT use this strategy when reaching without a cursor.

**Note:** During the localization task, instruct the participant to tap exactly above where they felt their right thumb crossed the arc (not where they think a cursor would have gone or anything strange like that).

#### For IMPLICIT version only: Explaining Perturbations/Rotation task

For these next few reaching tasks, the green cursor will move a bit differently, and you will need to compensate for this.

However you compensate, keep that strategy in mind since you will be asked to use this strategy several times, including when reaching without a cursor. Sometimes you will be asked to NOT use this strategy when reaching without a cursor.

#### WITHOUT Strategy – Exclusive

For this next task you will be reaching to a target without a cursor. For THESE trials, do not make use of any strategies you learned earlier and treat this as you did the original baseline reach-to-target task.

#### WITH Strategy - Inclusive

For this next task you will be reaching to a target without a cursor. For THESE trials, please make use of the strategy you learned earlier to correct for odd movement of the cursor.

### Appendix B: Clock-face image used to instruct participants on the nature of the perturbation. An animation was also provided

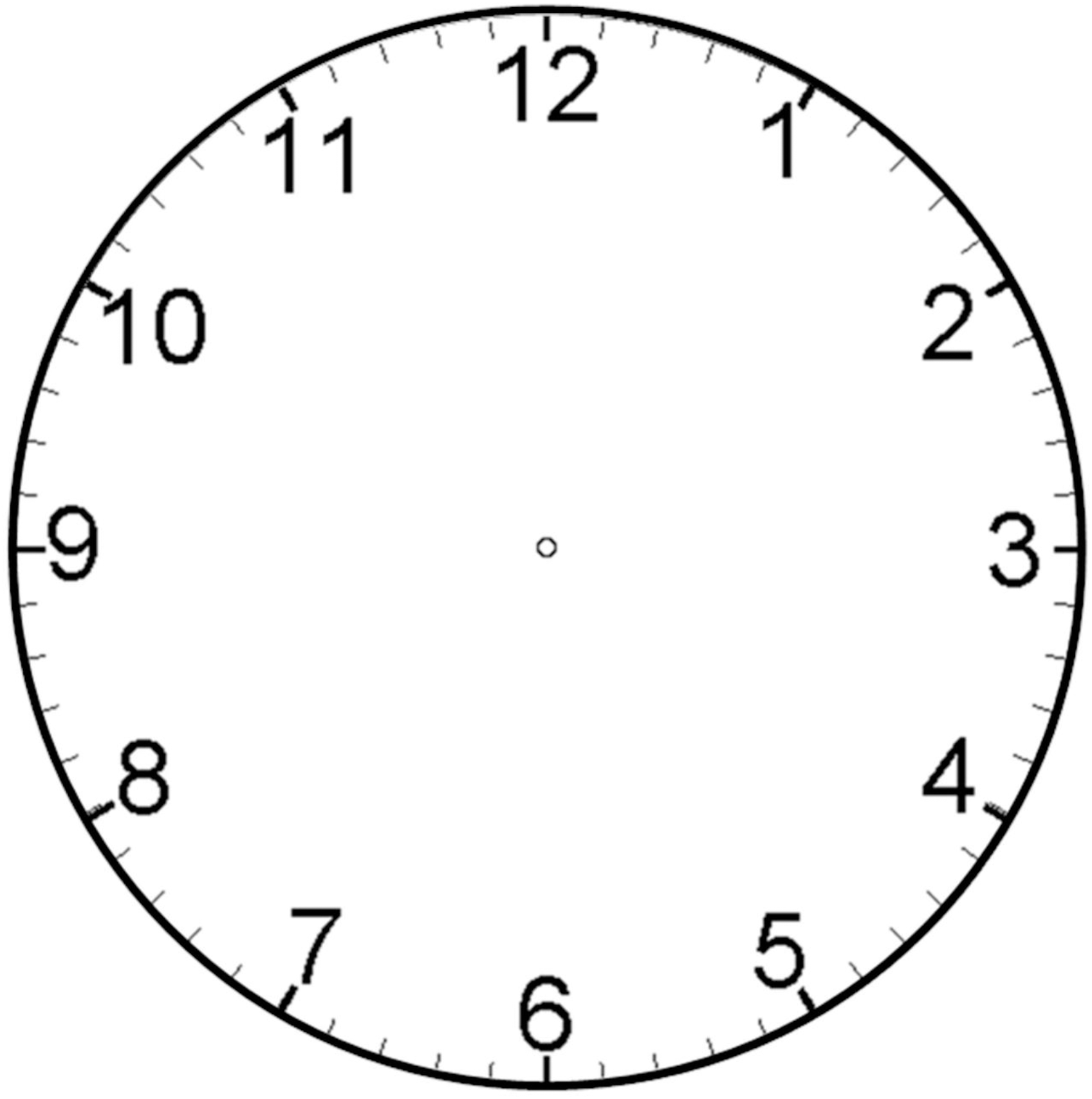

